# Homeostatic Turnover, Repair, and the Demographic Form of Haldane’s Principle

**DOI:** 10.64898/2026.09.20.753009

**Authors:** Ben Hardisty

## Abstract

Haldane’s principle relates equilibrium mutation load to deleterious mutational input rather than to selection strength. We study this balance in a homeostatically regulated, damage-structured population where mutation occurs during reproduction, repair restores individuals toward lower-damage states, and nonlinear demographic feedback maintains constant population size. An exact demographic identity shows that load at every time equals excess reproductive effort weighted by mean relative fecundity. Without repair, the equilibrium branch with positive pristine-class frequency is exactly solvable, and along it the load is exactly the mutation probability times the realized per-capita reproductive turnover rate; when the terminal cap is removed, this is precisely the realized deleterious input, for every mutation probability below one. For rare mutation it reduces to the mutation probability times the pristine turnover rate, with the local fitness contrast canceling at leading order. With a finite terminal class the branch instead reaches a closure-dependent boundary collision. Repair modifies the rare-mutation balance by partitioning first-order removal from the first damaged class between selection and restoration, multiplying the leading load by the selective share of that removal. An exact binary solution extends these results to finite mutation pressure: the unrepaired interior equilibrium collides nonhyperbolically with the fully damaged boundary and converges algebraically at the threshold, whereas any positive repair rate restores a unique interior equilibrium for positive mutation; at the no-repair threshold, its displacement from the boundary scales as the square root of the repair rate as repair vanishes. These results separate the familiar competing-transition structure of mutation–selection balance from the specifically homeostatic effects of turnover-dependent mutation input and nonlinear demographic feedback.

## 1. Introduction

Mutation–selection balance is one of the central organizing ideas of population genetics. Deleterious variants are continually introduced by mutation and removed by selection, and at equilibrium these opposing processes determine both their frequencies and their contribution to population fitness. In its classical form, Haldane’s principle states that the equilibrium mutation load is approximately equal to the deleterious mutation rate and, to leading order, is independent of the strength of selection against individual mutations [1, 2]. Stronger selection reduces the equilibrium frequency of a deleterious state, but each remaining individual carries a larger fitness deficit, causing the leading dependence on selection strength to cancel.

The scope of this cancellation is substantially broader than Haldane’s original setting. Hopf, Michod, and Sanderson [3] established Haldane-type load relations across a variety of reproductive systems. Kimura and Maruyama [4] showed that the epistatic reduction of mutation load under random mating does not occur under asexual reproduction, the setting adopted here. Bürger and Hofbauer [5] extended Haldane-type load results to arbitrary mutation patterns among alleles and to quantitative genetic traits, deriving first-order approximations for equilibrium mean fitness and mutation load; see also Bürger [6]. Hermisson et al. [7] developed a general deterministic haploid framework for mutation–selection balance, relating mutation load to ancestral structure and mean fitness and analyzing mutation-threshold phenomena. Together, these results show that neither selection-strength cancellation, backward transitions, nor mutation thresholds are confined to elementary one-locus models. Related threshold phenomena have also been studied extensively in quasispecies theory, where sufficiently strong mutation can delocalize a favored genotype [8, 9, 10]. The finite-*U* boundary transition derived below is distinct: total abundance is fixed by construction, and the transition is a collision, in class composition, of the interior equilibrium with the fully damaged boundary. For the finite multiclass hierarchy, Section 3.3 further shows that the corresponding collision is a consequence of the terminal closure rather than a threshold of the uncapped hierarchy.

A separate line of work has emphasized that mutation load must also be interpreted demographically. Steinsaltz, Evans, and Wachter [11] developed a nonlinear mutation–selection framework capable of incorporating full age-specific demographic schedules. Wachter, Evans, and Steinsaltz [12] subsequently obtained an exact nonlinear generalization of Haldane’s principle within this line of work. In this demographic setting, compensatory changes in fertility and survival can maintain near-zero long-term population growth despite genetic deterioration. Ecological treatments reach a related conclusion from a different direction: whether deleterious mutations reduce population performance depends on how density dependence and competition translate individual fitness differences into demographic consequences [13, 14]. That excess reproductive production can buffer genetic deaths without reducing population abundance is a longstanding qualitative observation [15]; the identity derived below gives this idea an exact dynamical form. Models in which population size itself responds to mutation load can instead generate feedback between genetic deterioration and demographic decline, including extinction [16]. More recently, a regulated two-type mutation–selection model has coupled mutation and selection directly to density-dependent population growth and yielded persistence and composition thresholds [17].

The present model addresses a different demographic question. Rather than allowing population size to respond to mutation load, we impose exact homeostasis: at every time, reproduction is rescaled by a state-dependent feedback variable so that realized reproduction balances current mortality. Mutation occurs during reproduction and is therefore multiplied by this same feedback, whereas mortality and repair are not. Writing

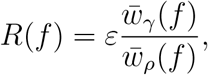

where 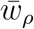 and 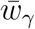 denote the population-mean relative fecundity and mortality, the reproductive and upward-mutational fluxes depend on the entire current class distribution. Thus the demographic cost of deleterious mutation is expressed not through declining abundance but through the additional reproductive effort required to maintain constant abundance.

This distinction also separates the model from standard fixed-operator mutation–selection systems such as that of Hermisson et al. [7]: here reproductive and upward-mutational fluxes depend on the current population through *R*(**f**), so the nonlinearity survives passage to class frequencies. This state-dependent compensation is the source of the load identities derived below; we return to the operator-level comparison in Section 4.

Repair introduces a second distinction. At the level of class transitions, restoration toward a less damaged state resembles backward mutation, and the competing effects of forward and backward transitions have a long history in mutation–selection theory [5, 7]. Here, however, repair is modeled as a conservative within-population transition occurring continuously in time, independently of reproduction. It changes the state occupied by an individual without itself producing an offspring or causing a death. Consequently, mutation input is tied to the homeostatically regulated reproductive flux, whereas repair provides a separate route by which damaged-state occupancy can be removed. The question is therefore not whether backward transitions can modify mutation–selection balance, but how an independent restoration pathway modifies the demographic expression of Haldane’s principle under exact homeostasis.

This formulation is relevant to populations in which abundance is tightly regulated while damage is continually generated and removed. Examples include stem-cell niches, stationary or resource-limited microbial populations, and other somatic compartments in which turnover replaces lost individuals while molecular or physiological repair can restore damaged states. Molecular lesions may be repaired, epigenetic or physiological states may be restored, and cellular damage may be partly reversed before selective loss of the affected individual [18, 19, 20]. Because mutation is coupled to reproduction whereas repair proceeds independently in continuous time, their relative effects depend on demographic turnover as well as on the fitness deficit associated with damage.

The contribution of the present model is not homeostatic regulation, backward transitions, or mutation thresholds individually, but the consequences of combining state-dependent exact homeostatic compensation with reproduction-linked mutation and independent conservative repair. This structure yields an exact off-equilibrium demographic identity linking mutation load directly to the excess reproductive effort required to maintain constant abundance. Under this structure, *εU* is the leading realized deleterious input per unit dimensionless time, so demographic turnover converts mutation probability per reproductive event into a realized input rate. In the exactly solvable binary system, any positive repair rate removes the no-repair boundary collision, while small repair displaces the threshold equilibrium from the fully damaged state on a square-root scale.

We formalize these processes in a haploid asexual population structured by damage class *i* = 0, 1, …, *n*. Reproduction and mortality vary geometrically across classes. Mutation moves offspring toward higher damage, repair moves individuals toward lower damage, and the state-dependent feedback *R*(**f**) maintains total population size exactly.

The analysis yields a sequence of structural results. First, mutation load has the exact demographic representation

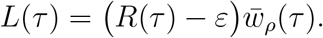

This identity holds for every biological state, not only at equilibrium, and its form contains no explicit repair term. It identifies mutation load with excess reproductive effort, weighted by mean relative fecundity, and therefore gives a direct demographic interpretation of load under exact homeostasis.

Second, in the absence of repair the multiclass equilibrium branch with positive pristine-class frequency is exactly solvable. Along this branch,

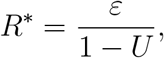

and the class frequencies satisfy explicit multiplicative recursions. For the finite terminal-class formulation used here, the branch exists precisely for

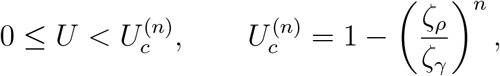

and approaches the fully damaged terminal state as 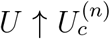. By contrast, when the terminal cap is removed, the corresponding pristine-positive equilibrium is normalizable for every *U <* 1; accordingly, 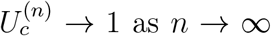.

For rare mutation,

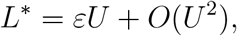

so demographic turnover sets the realized deleterious input per unit dimensionless time while the leading dependence on the fitness landscape cancels.

Third, when repair is present, the rare-mutation equilibrium branch obeys

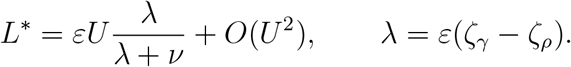

The factor *λ/*(*λ* + *ν*) is the leading-order share of removal from the first damaged class attributable to selection rather than repair. The competing-exit structure itself is familiar from mutation–selection systems with reverse transitions; the homeostatic result is that the realized input rate is *εU*, while demographic turnover enters the selective scale through *λ* = *ε*(*ζ*_*γ*_ − *ζ*_*ρ*_).

Finally, the binary specialization can be solved exactly at finite mutation pressure. Without repair, the general multiclass branch endpoint reduces to

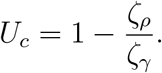

At this boundary the interior equilibrium reaches the fully damaged state; for *U* ≥ *U*_*c*_, the transition is one of class composition rather than population extinction, because total abundance remains homeostatically fixed.

Every positive repair rate instead yields a unique globally attracting interior equilibrium for *U >* 0. At the no-repair collision, convergence to the fully damaged boundary is algebraic, whereas small positive repair displaces the equilibrium from that boundary on a square-root scale. The binary solution therefore shows nonperturbatively how repair regularizes the boundary transition, while the multiclass rare-mutation result identifies the corresponding leading-order load mechanism.

## 2. Model

### 2.1. Damage structure and vital rates

We consider a haploid asexual population divided among damage classes

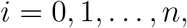

for a fixed integer *n* ≥ 1, where *i* = 0 is the pristine class and increasing *i* represents increasing damage burden. Let *N*_*i*_(*t*) denote the number of individuals in class *i*, let

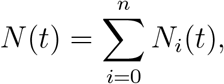

and define the class frequencies

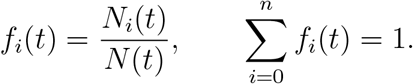

Write

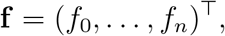

and denote the probability simplex by

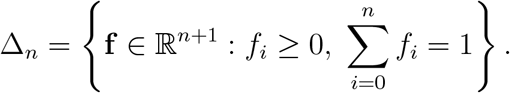

For 0 ≤ *i* ≤ *n*, let **e**_*i*_ denote the vertex of Δ_*n*_ concentrated entirely in damage class *i*.

Class-specific reproduction and mortality rates are

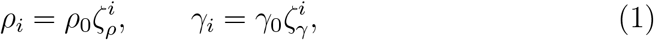

where

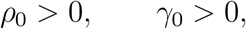

and

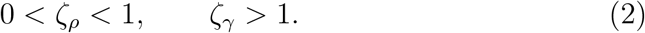

The geometric parameterization is adopted for concreteness; the principal load results extend to arbitrary relative rate sequences as described in Remark 1. Accumulated damage therefore reduces reproductive potential and increases mortality. We measure time in units of the pristine reproductive time,

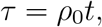

and define the dimensionless demographic ratio

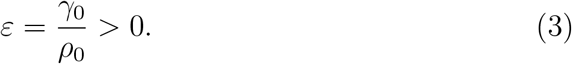

Values 0 *< ε <* 1, for which the pristine baseline reproductive rate exceeds the pristine mortality rate, are the primary biological regime of interest, but none of the results below requires this upper bound.

During reproduction, an offspring acquires one additional unit of damage with probability

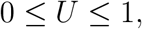

and otherwise retains its parent’s damage class. Repair occurs independently of reproduction: an individual in class *i* ≥ 1 moves to class *i* − 1 at dimensionless rate

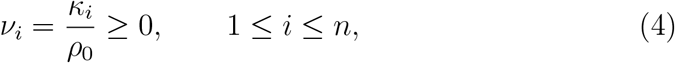

where *κ*_*i*_ is the corresponding dimensional repair rate and *ν*_0_ = 0.

Repair is a conservative state transition. It moves individuals between classes but does not itself create or remove individuals from the population.

The upper class *n* is treated as terminal. Reproduction by a class-*n* individual remains in class *n*, regardless of whether an additional damage event occurs. This convention closes the finite damage hierarchy without introducing flux out of the represented state space. Total population conservation follows from accounting for all birth, mutation, and repair destinations together with the homeostatic feedback. The terminal convention does not affect the leading rare-mutation load derived below, but it does determine the finite-*U* endpoint of the pristine-positive no-repair branch. Accordingly, that endpoint is a property of the finite capped hierarchy rather than a universal mutation threshold.

### 2.2. Homeostatic feedback

Total population size is maintained by a feedback variable that rescales all reproductive rates. Define the mean relative fecundity and mortality by

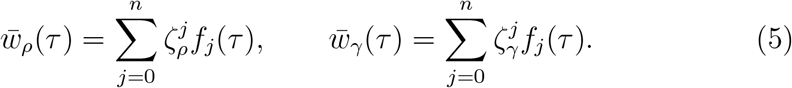

The homeostatic feedback ratio is

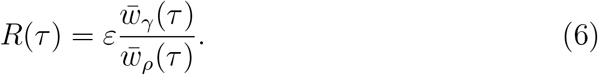

Equivalently, realized reproduction exactly balances mortality:

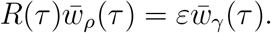

The feedback *R*(*τ*) is a positive reproductive multiplier and is not restricted to be at most one; the rates *ρ*_*i*_ therefore represent baseline class-specific reproductive rates rather than hard upper bounds on reproductive output. Consequently,

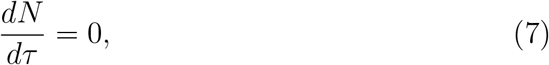

and the equations for class abundances and class frequencies coincide up to the constant factor *N*.

The feedback *R*(*τ*) is defined algebraically by the current class distribution. It is not a quasi-steady approximation and no fast–slow reduction is imposed.

### 2.3. Multiclass damage–repair dynamics

The pristine class satisfies

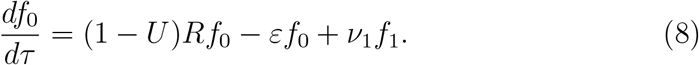

For interior classes 1 ≤ *i < n*,

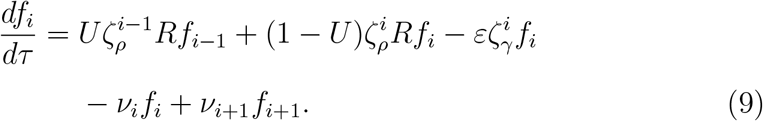

The terminal class obeys

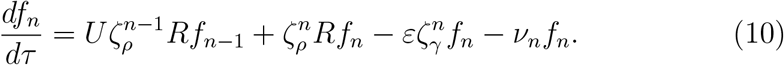

The absence of a factor 1 − *U* in the terminal self-renewal term reflects the fact that all class-*n* offspring remain in the terminal class.

Summing Eqs. (8)–(10) causes all mutation fluxes and repair fluxes to cancel pairwise. The remaining birth and death terms cancel by Eq. (6), giving

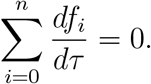

The probability simplex Δ_*n*_ is forward invariant. Indeed,

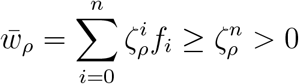

on Δ_*n*_, so *R*(**f**) is finite and positive. On a boundary face where *f*_*i*_ = 0, the corresponding derivative is

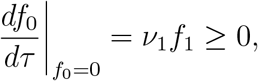

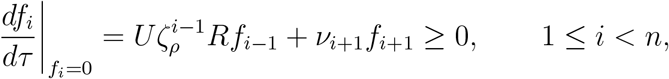

and

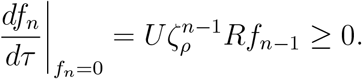

Together with population conservation, these inequalities show that solutions beginning in Δ_*n*_ remain in Δ_*n*_.

Repair therefore affects homeostatic feedback only indirectly, through its effect on the class distribution.

### 2.4. Mutation load and its demographic representation

At the mutation-free pristine equilibrium,

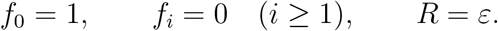

Because exact homeostasis enforces zero realized population growth at all times, mutation load cannot be identified here with a reduction in realized mean population growth. We therefore measure the composition-weighted demographic deficit relative to the mutation-free pristine environment. This pristine reference environment is fixed by construction and is not reevaluated at the current feedback *R*(**f**). In that environment, the net growth-rate deficit of class *i* relative to the pristine class is

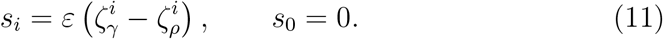

Because repair is modeled as a conservative transition, it does not enter *s*_*i*_. Repair alters the time spent in a damaged class, but not that class’s intrinsic reproductive or mortality deficit.

We define the instantaneous mutation load by

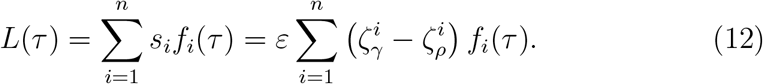

At an equilibrium 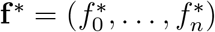, we write

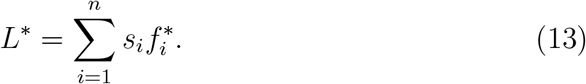

For comparison with the conventional unit-coefficient form of Haldane’s principle, it is useful to normalize the demographic load by the pristine homeostatic turnover scale:

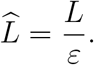

### 2.5. Exact binary specialization

To obtain a nonperturbative repair result, we specialize the model to one pristine class and one terminal damaged class. Set *n* = 1 and define

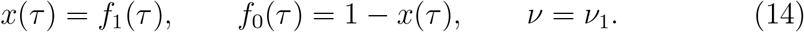

The two class equations are

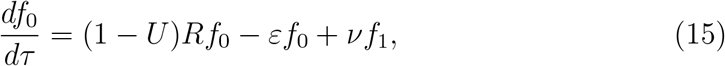

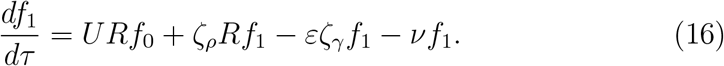

Because class 1 is terminal, its full reproductive output remains in class 1.

For later convenience, define

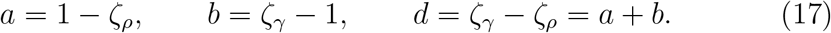

Since *f*_0_ = 1 − *x*, the feedback becomes

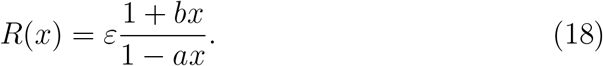

Substitution into Eq. (16) reduces the full nonlinear homeostatic system to the scalar equation

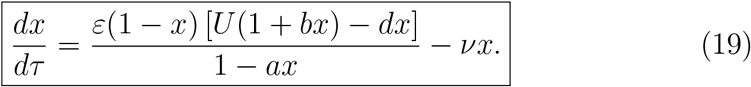

The biological state space 0 ≤ *x* ≤ 1 is forward invariant. At *x* = 0,

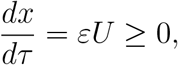

and at *x* = 1,

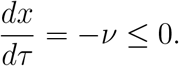

At equilibrium, Eq. (19) gives

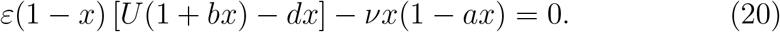

Define

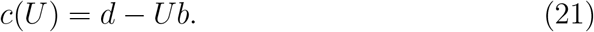

Then the equilibrium equation is the quadratic

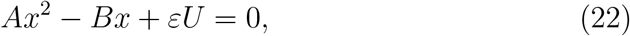

where

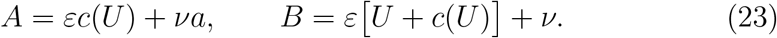

The biologically admissible root and its stability are derived below.

For the binary model, the damaged-state selection coefficient is

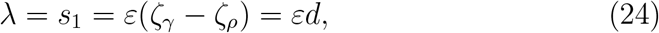

and the load reduces exactly to

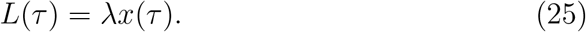

The exact finite-*U* equilibrium load is therefore obtained directly from the admissible root of Eq. (22).

## 3. Results

The results proceed from general structural properties to exact special cases. We first establish a demographic identity that holds for every biological state and then characterize the analytic rare-mutation equilibrium branch issuing from the pristine state. We next derive the exact no-repair benchmark and the repair-modified Haldane principle for the general multiclass model. Finally, we solve the binary damage–repair model exactly for arbitrary mutation and repair rates.

### 3.1. An exact demographic identity

#### Proposition 1

(Exact demographic representation of load). *For every biological state* **f** ∈ Δ_*n*_ *and for arbitrary conservative repair rates*,

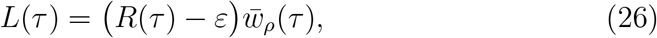

*where*

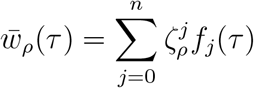

*is the population-mean fecundity relative to the pristine class. Proof*. By the definition of the homeostatic feedback,

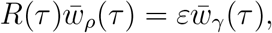

where

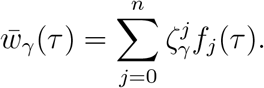

Subtracting 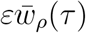 from both sides gives

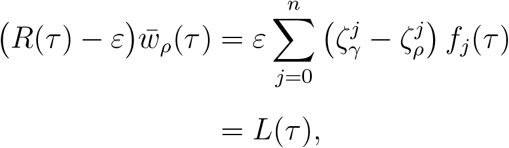

by Eq. (12).

Proposition 1 gives the mutation load a direct demographic interpretation. The quantity *R* − *ε* is the reproductive effort in excess of the pristine mortality-matched baseline, and 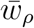 weights that effort by the population’s mean relative fecundity. The identity is an immediate algebraic consequence of the definition of *R*; its significance is interpretive rather than technical, since it holds away from equilibrium as well as at equilibrium and identifies load directly with excess reproductive effort. Its form contains no explicit repair term, although repair affects the class distribution and therefore the values of *L, R*, and 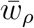.

In terms of the turnover-normalized load introduced in Section 2.4, Proposition 1 becomes

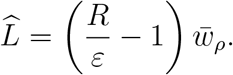

The quantity 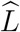 is a turnover-normalized growth-rate deficit, not a fraction of mean fitness, and need not lie between zero and one.

*Two load conventions*. It is also useful to relate the fixed-reference load *L* to the corresponding load defined in the realized homeostatic environment. Define the realized per-capita growth rate of class *i* by

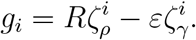

By Eq. (6),

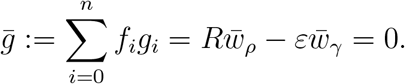

Moreover, *g*_0_ = *R* − *ε*, and for every *i* ≥ 1,

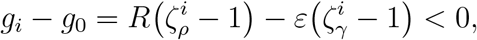

so the pristine class has the maximal realized growth rate. Thus the usual maximum-minus-mean load in realized growth-rate units is

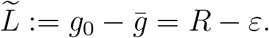

Proposition 1 therefore gives the exact relation

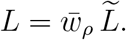

Hence load measured in the fixed pristine reference environment and load measured in the realized homeostatic environment differ exactly by the population’s mean relative fecundity. Along the rare-mutation branch, 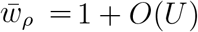 and 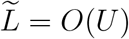, so

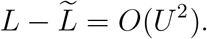

The two conventions therefore give the same leading-order Haldane relation, although they differ at finite mutation pressure. We retain *L* because its coefficients *s*_*i*_ are fixed parameters of the model rather than functions of the current population state.

### 3.2. Rare-mutation equilibrium branch

The equilibrium branch issuing from the pristine state has a strict mutation-order hierarchy: reaching damage class *i* requires at least *i* upward mutation events, even though repair couples higher classes back into lower classes.

#### Proposition 2

(Rare-mutation branch and mutation-order hierarchy). *Fix* 1 ≤ *n <* ∞, *ε >* 0, 0 *< ζ*_*ρ*_ *<* 1 *< ζ*_*γ*_, *and finite repair rates ν*_*i*_ ≥ 0 *that are independent of U*. *Define*

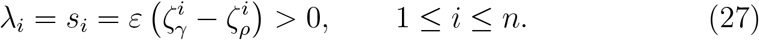

*There is a locally unique equilibrium branch*

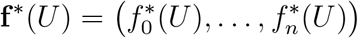

*analytic in a neighborhood of U* = 0 *and satisfying* **f** ^∗^(0) = **e**_0_. *Along this branch*,

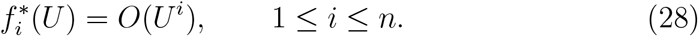

*Equivalently*,

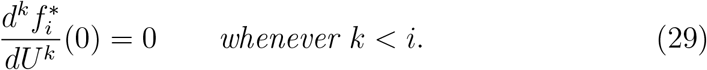

*More precisely*,

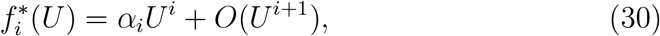

*where*

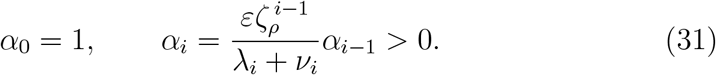

*In particular*,

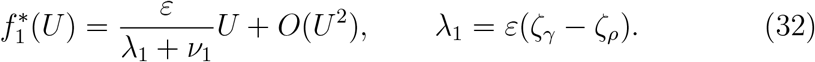

*Consequently, the branch lies in the biological interior for all sufficiently small U >* 0. *Moreover, this branch is locally asymptotically stable within* Δ_*n*_ *for all sufficiently small U >* 0.

*Proof*. On the simplex, use

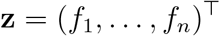

as independent variables and write

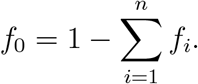

Let

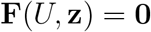

denote the damaged-class equilibrium equations. Since 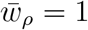 at the pristine state and remains positive in a neighborhood of that state, the feedback *R*(**f**), and hence **F**, is analytic near (*U*, **z**) = (0, **0**).

At the pristine equilibrium, the Jacobian

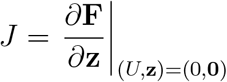

is upper bidiagonal, with

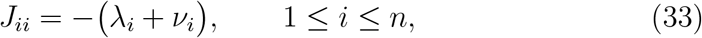

and

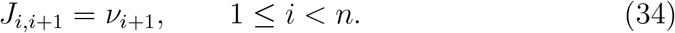

All remaining entries vanish. Since every diagonal entry is strictly negative, *J* is invertible. The implicit function theorem therefore gives a locally unique equilibrium branch **z**^∗^(*U*) analytic near *U* = 0. Because *J* is upper bidiagonal, its spectrum is 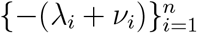, which lies strictly in the left half-plane. The pristine equilibrium is therefore hyperbolically stable within the simplex, and by continuity of the spectrum the branch **f** ^∗^(*U*) is locally asymptotically stable within Δ_*n*_ for all sufficiently small *U >* 0.

Expand

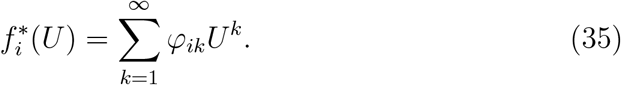

We prove by induction on *k* that

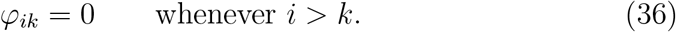

At order *U*, the only direct source from the pristine state is the mutational flux into class 1. Consequently, the coefficient vector

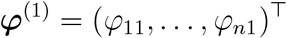

satisfies

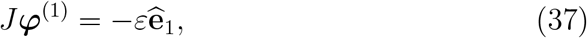

where 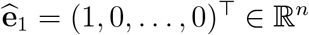. Backward substitution gives

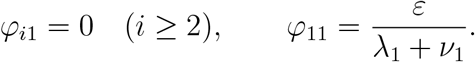

Now suppose that

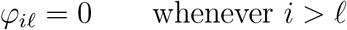

for every *l < k*. At order *U*^*k*^, the equilibrium equations can be written as

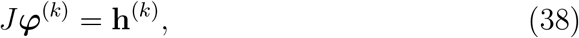

where **h**^(*k*)^ depends only on coefficients of orders strictly less than *k*.

For class *i*, the only term that creates occupancy from a lower damage class is

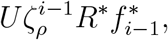

where *R*^∗^ = *R*(**f** ^∗^(*U*)). Its coefficient at order *U*^*k*^ depends only on coefficients of 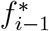 of orders at most *k* − 1. When *i > k*, all such coefficients vanish by the induction hypothesis.

After the linear contribution *J φ*^(*k*)^ is separated, the remaining order-*U*^*k*^ terms in the class-*i* equation involve only coefficients of 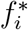 of orders strictly less than *k*, multiplied by positive-order terms arising from *R*^∗^ − *ε* or, for *i < n*, from the factor 1 − *U*. These coefficients also vanish when *i > k* by the induction hypothesis. Hence

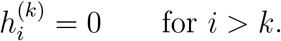

Because *J* is upper triangular, backward substitution in Eq. (38) gives

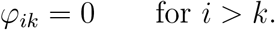

Induction over *k* proves Eq. (28).

It remains to identify the first nonzero coefficient of each class. At order *U*^*i*^, repair inflow from class *i* + 1 is absent because 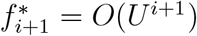, and

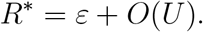

The order-*U*^*i*^ balance in the class-*i* equation is therefore

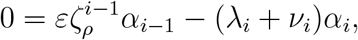

which gives Eq. (31). Since *α*_0_ = 1 and all factors in the recursion are positive, *α*_*i*_ *>* 0 for every *i*. Thus 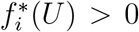 for sufficiently small *U >* 0, while

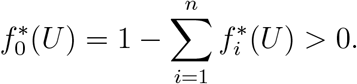

The branch is therefore biologically admissible near *U* = 0.

Proposition 2 implies

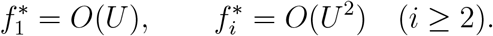

Hence only the first damaged class contributes to mutation load at order *U*. Repair rates *ν*_*i*_ for *i* ≥ 2 enter only through the *O*(*U* ^2^) remainder, whereas

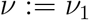

controls the leading repair correction.

Along this branch,

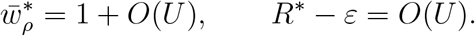

Proposition 1 therefore gives

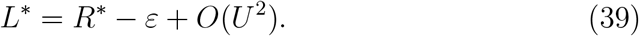

Thus the leading-order equality between mutation load and excess reproductive effort is the rare-mutation limit of the exact demographic identity (26).

### 3.3. Exact no-repair benchmark

#### Proposition 3

(Exact no-repair benchmark and branch domain). *Set*

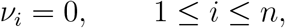

*and define*

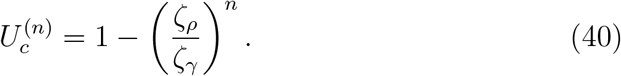

*The no-repair system has an equilibrium satisfying* 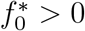 *if and only if*

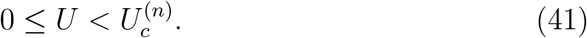

*Within this range, the equilibrium with* 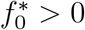 *is unique*.

*Along this branch, the homeostatic feedback is*

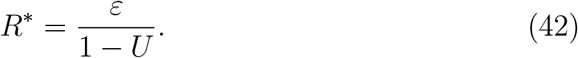

*For the nonterminal classes* 1 ≤ *i < n*,

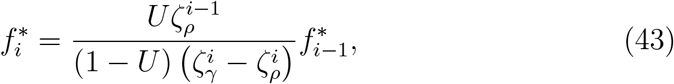

*whereas the terminal class satisfies*

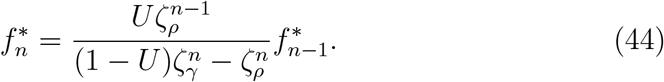

*As the mutation probability approaches the branch endpoint*,

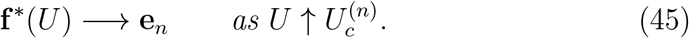

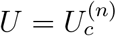, *no equilibrium with* 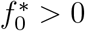 *remains*.

*Moreover, along the pristine-positive branch the load satisfies the exact relation*

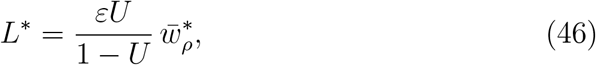

*and hence, for rare mutation*,

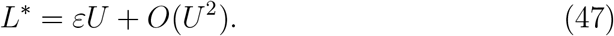

*Thus the leading-order load is independent of ζ*_*ρ*_ *and ζ*_*γ*_.

*Proof*. With repair absent, the equilibrium equation for the pristine class is

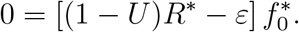

Because 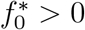,

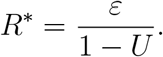

In particular, any equilibrium with 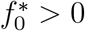 must satisfy *U <* 1.

For 1 ≤ *i < n*, the equilibrium equation is

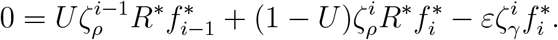

Substitution of Eq. (42) and rearrangement give Eq. (43).

For the terminal class,

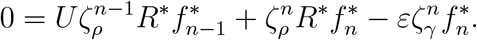

Substituting Eq. (42) gives

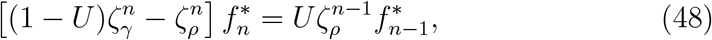

which is equivalent to Eq. (44) whenever the coefficient on the left is nonzero.

For 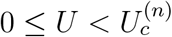,define

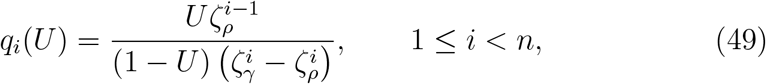

and

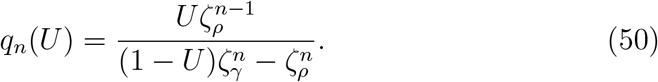

The denominator in Eq. (50) is positive precisely when

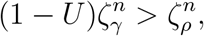

or equivalently,

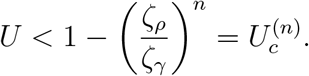

Let

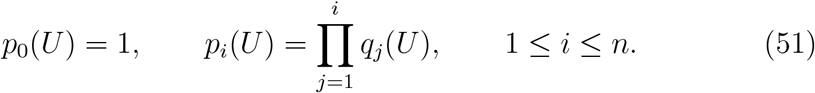

Successive application of the equilibrium recursions gives

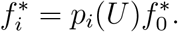

Normalization therefore yields

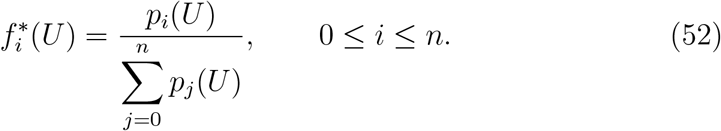

All components are nonnegative and 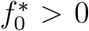. Because the recursions determine every 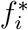 as a multiple of 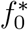, and normalization then determines 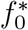, this equilibrium is unique among equilibria satisfying 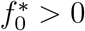.

It remains to show that the stated domain is necessary. Suppose 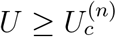 and that an equilibrium with 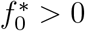 exists. By the preceding argument, such an equilibrium must also satisfy *U <* 1. Since 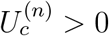, we have *U >* 0, and the interior recursions imply successively that

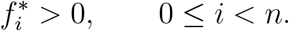

For *n* = 1, this statement reduces simply to the assumed inequality 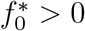.

At 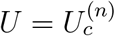, the coefficient on the left side of Eq. (48) vanishes, whereas its right side is strictly positive. The terminal balance therefore cannot hold. For 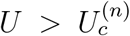, the coefficient on the left is negative while the right side remains positive, so Eq. (48) would require 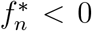. Hence no biologically admissible equilibrium with 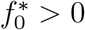 exists when 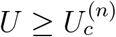.

Finally, as 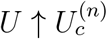, the quantities *q*_1_, …, *qn* _*−*1_ remain finite, whereas

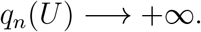

Consequently, *p*_0_, …, *p*_*n*−1_ remain finite while *p*_*n*_ → ∞. Equation (52) therefore gives

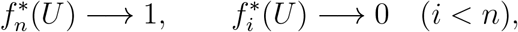

which proves Eq. (45).

To obtain the load, Proposition 1 and Eq. (42) give the exact relation

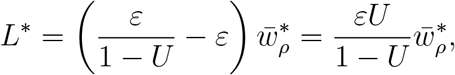

which proves Eq. (46). By Proposition 2, specialized to *ν*_*i*_ = 0,

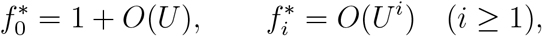

and hence

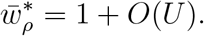

Therefore,

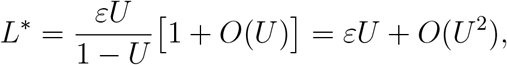

which proves Eq. (47).

Proposition 3 identifies the exact biological domain of the no-repair equilibrium branch issuing from the pristine state. It does not assert global attraction of that branch for arbitrary *n*. At 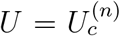, the branch reaches the fully damaged vertex **e**_*n*_, which is an equilibrium of the no-repair system for every *U*. Thus 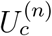 marks a boundary collision in class composition rather than population extinction.

*No-repair face equilibria*. The recursion in Proposition 3 also characterizes the remaining boundary equilibria of the finite no-repair hierarchy. Suppose an equilibrium has lowest occupied class *k < n*, so that

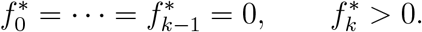

The class-*k* balance then gives

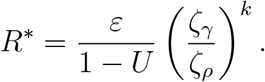

After shifting the class index by *k*, the higher-class balances reduce to the same recursions as in Proposition 3 for a hierarchy of length *n* − *k*. Hence this face branch exists precisely for

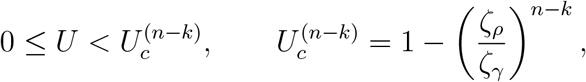

and approaches **e**_*n*_ at its endpoint.

For *U >* 0, positive occupancy of the lowest occupied class *k < n* forces positive occupancy successively in every class *j > k* through the upward-mutational recursions, so no gapped equilibrium support is possible. At *U* = 0, simultaneous occupancy of two distinct classes would require

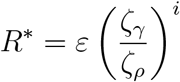

for two different values of *i*, which is impossible; hence the only *U* = 0 equilibria are the vertices **e**_*k*_. For 0 ≤ *k < n*, these vertices are precisely the *U* = 0 members of the face branches described above, while **e**_*n*_ is the terminal equilibrium. Together, these branches and **e**_*n*_, which exists for every *U*, exhaust the equilibrium set of the finite no-repair system. In particular, because 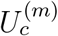 is increasing in *m*, for 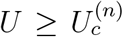no nonterminal face branch remains and **e**_*n*_ is the unique equilibrium.

Moreover, at **e**_*n*_ the reduced Jacobian is lower bidiagonal, with diagonal entries

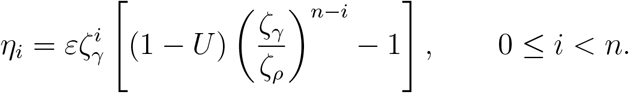

Its eigenvalues are therefore precisely 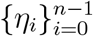. Thus **e**_*n*_ is locally asymptotically stable for 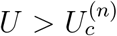, while at 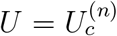 the eigenvalue associated with invasion of the pristine class vanishes, consistently with the nonhyperbolic boundary collision.

For 1 ≤ *k < n*, these face equilibria are transversely unstable to introduction of the immediately lower damage class, whose linear growth rate is

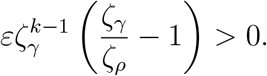

#### Proposition 4

(No finite threshold in the uncapped no-repair hierarchy). *Consider the no-repair model with damage classes i* = 0, 1, 2, …, *obtained by removing the terminal class and applying the nonterminal dynamics to every i* ≥ 1. *For the geometric rates in Eq*. (1), *there is a unique normalizable equilibrium with* 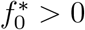 *for every* 0 ≤ *U <* 1.

*Along this equilibrium*,

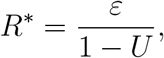

*and*

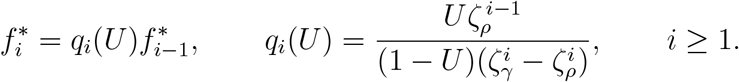

*Consequently, the finite-class endpoints*

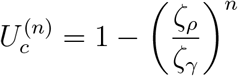

*satisfy* 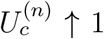 *as n* → ∞. *Thus the finite-U boundary collision of Proposition 3 is a consequence of the terminal closure: no corresponding pristine-positive threshold occurs below U* = 1 *in the uncapped hierarchy*.

*Proof*. If 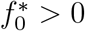, the pristine-class equilibrium equation gives

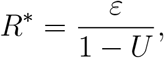

so necessarily *U <* 1. For fixed 0 ≤ *U <* 1, define

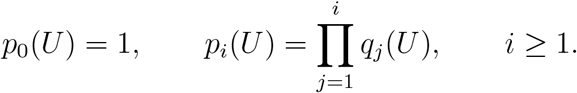

Writing

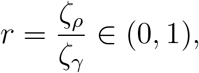

we have

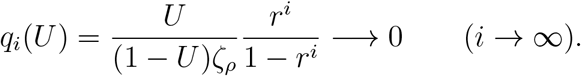

Hence

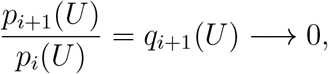

So

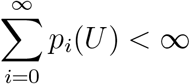

by the ratio test. Therefore

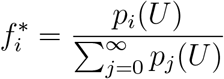

defines a probability distribution with 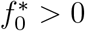.

The weighted sums defining the homeostatic feedback are also finite. In-deed,

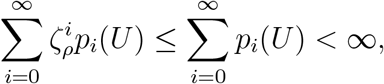

while

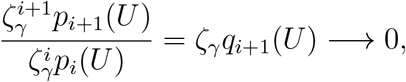

so

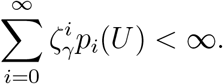

Since 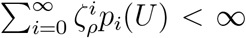 the boundary mutation flux tends to zero as the truncation index tends to infinity. The equilibrium recursions may therefore be summed over all classes, and their mutation fluxes telescope, giving

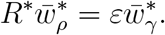

Thus the feedback *R*^∗^ = *ε/*(1 − *U*) is self-consistent.

Any equilibrium with 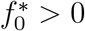 must satisfy the same pristine-class balance and the same successive recursions, so normalization makes this equilibrium unique. At *U* = 1, the pristine-class balance cannot hold with 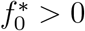. Hence the pristine-positive branch exists precisely for 0 ≤ *U <* 1.

#### Exact mutational-input identity

Along the pristine-positive no-repair branch, Eq. (42) and Proposition 1 give

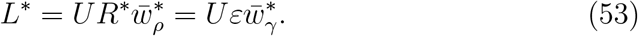

Thus the equilibrium load is exactly *U* times the realized per-capita reproductive turnover rate. In the finite capped hierarchy, the actual upward class-transition flux is

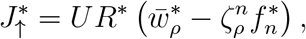

so that

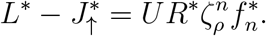

This discrepancy is entirely a consequence of the terminal closure. In the uncapped hierarchy of Proposition 4, every mutational event produces an upward transition, and therefore

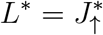

exactly for every 0 ≤ *U <* 1.

##### Remark 1

(Role of the geometric parameterization). The geometric form (1) is used for concreteness. Proposition 1, Proposition 2, Proposition 3, Eq. (53), and Corollary 1 hold verbatim for arbitrary relative rate sequences

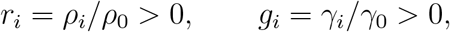

with *r*_0_ = *g*_0_ = 1 and *g*_*i*_ *> r*_*i*_ for *i* ≥ 1, on replacing 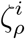 by *r*_*i*_ and 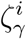 by *g*_*i*_ throughout. Then

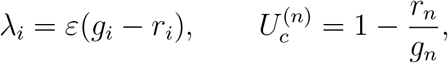

so the endpoint of the pristine-positive branch depends only on the relative rates of the terminal class.

Theorem 1 likewise holds with *ζ*_*ρ*_ and *ζ*_*γ*_ replaced by *r*_1_ and *g*_1_, provided 0 *< r*_1_ *<* 1 *< g*_1_. No monotonicity of the sequences *r*_*i*_ and *g*_*i*_ across damage classes is required. Thus the principal load results are not tied to geometric or multiplicative dependence of the vital rates on damage class.

The face-equilibrium classification following Proposition 3, by contrast, uses the geometric structure through the index shift, so the endpoints 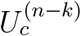 and the associated stability classification are not asserted here for arbitrary rate sequences. Likewise, Proposition 4 requires additional summability conditions for general sequences to ensure normalization and finiteness of the homeostatic feedback. Related existence questions on countably infinite trait spaces have recently been addressed by Pollett [21], whose operator-theoretic treatment of confining selection suggests a natural approach to studying such an extension; the comparison is developed in Section 4.

### 3.4. Repair-modified Haldane principle

#### Corollary 1

(Repair-modified Haldane principle). *Consider the rare-mutation equilibrium branch of the multiclass model. Fix*

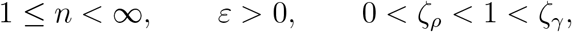

*and finite repair rates ν*_*i*_ ≥ 0 *that are independent of U*. *Write*

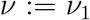

*for the repair rate from class* 1 *to class* 0. *Then*

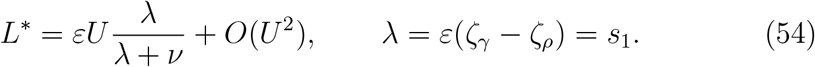

*Thus, at leading order, repair multiplies the unrepaired homeostatic load by the fraction of first-order damaged-state removal attributable to selection rather than repair*.

*Proof*. By Proposition 2,

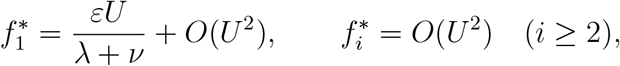

where

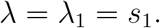

Therefore,

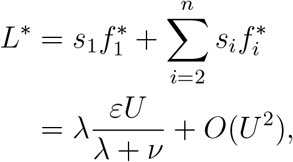

which proves Eq. (54).

Corollary 1 separates the leading load into two factors:

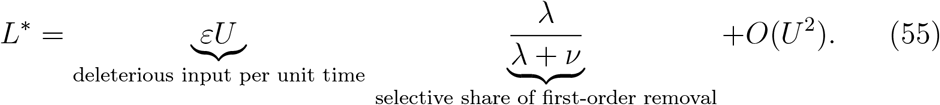

At the pristine homeostatic state, reproduction occurs at rate *ε*, so a mutation probability *U* per reproductive event produces deleterious input at rate *εU*. Once an individual enters the first damaged class, it is depleted from that class through selection at the effective leading rate *λ* or through repair at rate *ν*. The ratio *λ/*(*λ* + *ν*) is therefore the selective share of total first-order removal.

When *ν*= 0, the first-order repair pathway is absent and Eq. (47) is recovered. When *ν>* 0, repair reduces first-damaged-class occupancy before selection removes those individuals, thereby lowering the leading-order mutation load. At leading order, the fitness landscape enters only through the local class-0 to class-1 contrast *λ* = *s*_1_; higher-class coefficients begin to enter at order *U* ^2^, starting with *s*_2_.

Proposition 1 also gives the corresponding feedback expansion without requiring a separate corollary:

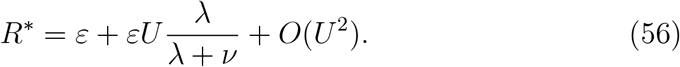

Thus repair suppresses the excess reproductive effort and the mutation load by the same leading-order factor.

Equivalently, under the turnover normalization,

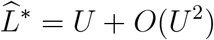

without repair, while with repair

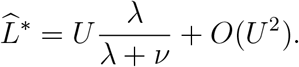

Thus, in the no-repair case, the conventional unit Haldane coefficient is recovered independently of the demographic ratio *ε*.

#### Remark 2

(Rare-mutation slope). Let

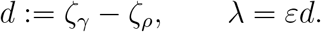

Corollary 1 gives

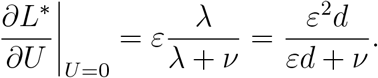

The demographic and repair limits of this slope are discussed in Section 4.

### 3.5. Exact binary equilibrium and repair regularization

Corollary 1 applies to an arbitrary number of damage classes but is asymptotic in *U*. The binary specialization introduced in Section 2.5 can instead be solved exactly for arbitrary mutation and repair rates, and doing so reveals a threshold phenomenon that the rare-mutation expansion cannot resolve.

#### Theorem 1

(Exact binary equilibrium and repair regularization). *Consider the binary model* (15)*–*(16), *with*

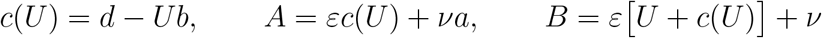

*as in* Eqs. (21) *and* (23), *and let*

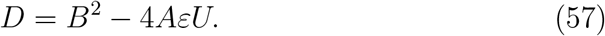

*Assume ε >* 0 *as above*.

i. (Repair present.) *If* 0 *< U* ≤ 1 *and ν>* 0, *the system has a unique equilibrium in the biological interior* 0 *< x <* 1, *given by*

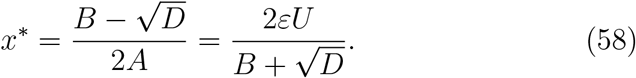

*This equilibrium is globally attracting on* 0 ≤ *x* ≤ 1. *Its exact homeostatic feedback and mutation load are*

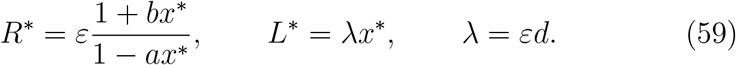

*For fixed U >* 0, *both x*^∗^ *and L*^∗^ *are strictly decreasing functions of the repair rate ν. As U* → 0,

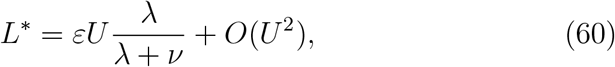

*recovering Corollary 1*. *When U* = 0 *and ν>* 0, *the unique attracting equilibrium is the pristine boundary state x*^∗^ = 0.
ii. (Repair absent.) *If ν*= 0, *define*

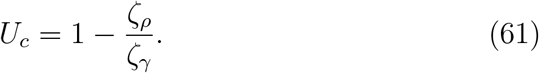

*When U* = 0, *the pristine equilibrium x*^∗^ = 0 *attracts every solution with* 0 ≤ *x*(0) *<* 1, *while the fully damaged boundary state x* = 1 *is an unstable equilibrium. For* 0 *< U < U*_*c*_, *the unique attracting interior equilibrium is*

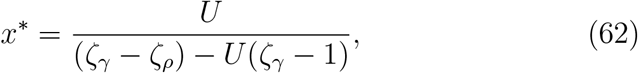

*with exact load*

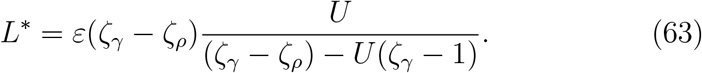

*At U* = *U*_*c*_, *the interior equilibrium reaches the fully damaged boundary x* = 1. *For U* ≥ *U*_*c*_, *the attracting biological equilibrium is*

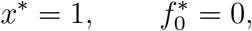

*and the load saturates at the full damaged-state deficit*,

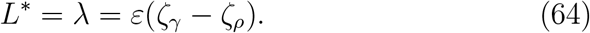

*Consequently, for every* 0 *< U* ≤ 1, *any positive repair rate removes the no-repair boundary transition of part (ii) and restores the unique interior equilibrium of part (i)*.

*Proof. Part (i)*. For 0 *< U* ≤ 1,

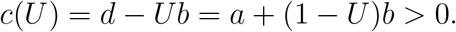

Consequently,

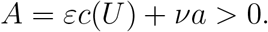

Multiplying the equilibrium form of Eq. (19) by 1 − *ax*, which satisfies

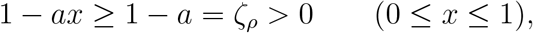

Gives

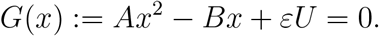

At the boundaries,

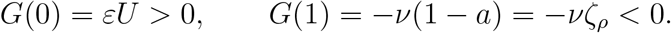

Hence *G* has a root in (0, 1). Since *A >* 0, the quadratic opens upward; because *G*(1) *<* 0, its second root lies above 1. The two roots are therefore distinct, so *D >* 0, and the unique biologically admissible root is the smaller root (58). The rationalized expression in Eq. (58) is algebraically equivalent and is numerically stable when *U* is small.

Because 1 − *ax >* 0 on the biological interval, the scalar vector field has the same sign as *G*(*x*):

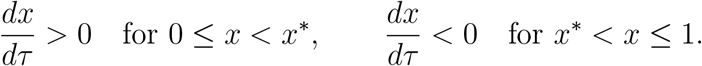

Every solution beginning below *x*^∗^ therefore increases monotonically toward *x*^∗^, while every solution beginning above *x*^∗^ decreases monotonically toward it. Thus *x*^∗^ is globally attracting on the biological interval. Equation (18) gives the exact feedback and Eq. (25) gives *L*^∗^ = *λx*^∗^.

The dependence on repair rate is also monotone. Since

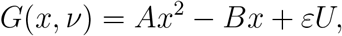

we have

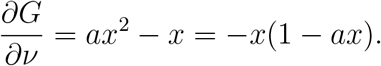

At the biologically admissible smaller root,

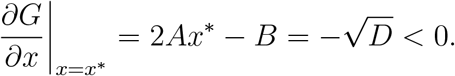

Implicit differentiation therefore gives

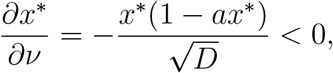

because 0 *< x*^∗^ *<* 1 and 1 − *ax*^∗^ *>* 0. Since *L*^∗^ = *λx*^∗^ with *λ >* 0,

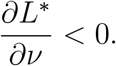

Finally, as *U* → 0,

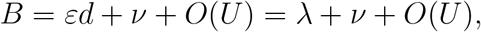

and

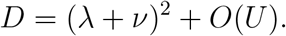

Since *λ* + *ν>* 0,

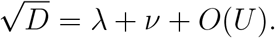

Using the rationalized form of Eq. (58) therefore gives

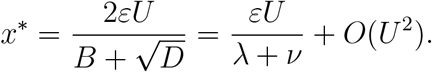

Multiplication by *λ* gives Eq. (60).

*Part (ii)*. When *ν*= 0, the equilibrium equation (20) factors as

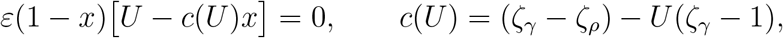

so the algebraic equilibria are *x* = 1 and *x* = *U/c*(*U*). Since *c*(*U*) is linear and decreasing in *U* with *c*(0) = *ζ*_*γ*_ − *ζ*_*ρ*_ *>* 0 and *c*(1) = 1 − *ζ*_*ρ*_ *>* 0, we have *c*(*U*) *>* 0 throughout 0 ≤ *U* ≤ 1. The second equilibrium lies in (0, 1) exactly when *U < c*(*U*), equivalently *Uζ*_*γ*_ *< ζ*_*γ*_ − *ζ*_*ρ*_, which is precisely *U < U*_*c*_ with *U*_*c*_ given by Eq. (61).

When *U* = 0, Eq. (19) reduces to

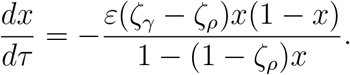

The denominator is positive on 0 ≤ *x* ≤ 1, so the vector field is strictly negative for 0 *< x <* 1. Hence every solution with 0 ≤ *x*(0) *<* 1 converges monotonically to *x* = 0, while *x* = 1 is an unstable boundary equilibrium.

For 0 *< U < U*_*c*_ the vector field is positive below the interior root and negative above it, so that root is attracting; it is Eq. (62), and Eq. (25) gives Eq. (63).

At *U* = *U*_*c*_ we have *c*(*U*_*c*_) = *U*_*c*_, and the scalar equation reduces to

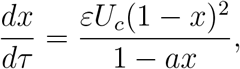

which is positive for 0 ≤ *x <* 1. Thus the two equilibria have merged at *x* = 1, which is attracting from within the biological interval.

At the collision point the boundary equilibrium is nonhyperbolic. Writing

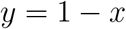

Gives

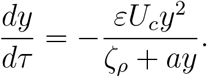

Hence, as *y* → 0,

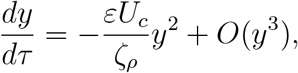

and therefore

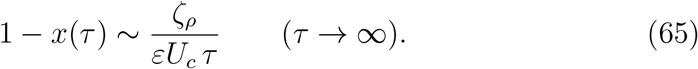

Thus convergence to the fully damaged boundary is algebraic at the threshold rather than exponential. For *U > U*_*c*_ the second algebraic root exceeds 1 and the vector field is positive throughout 0 ≤ *x <* 1, so *x* = 1 is again attracting. At and above the threshold, Eq. (25) gives *L*^∗^ = *λx*^∗^ = *λ*, which is Eq. (64). For every 0 *< U* ≤ 1, Part (i) shows that any *ν>* 0 instead yields a unique equilibrium strictly between 0 and 1.

#### Remark 3

(Zero-repair limit and threshold regularization). For fixed *U >* 0, the repaired equilibrium satisfies

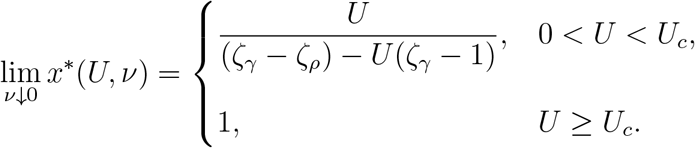

Thus the positive-repair equilibrium converges to the interior no-repair branch below *U*_*c*_ and to the fully damaged boundary at and above *U*_*c*_.

The approach is singular at the collision point. At *U* = *U*_*c*_, writing

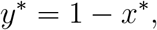

the equilibrium condition becomes

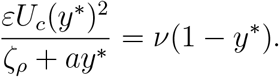

Since *y*^∗^ → 0 as *ν*↓ 0, it follows that

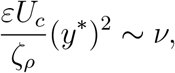

and hence

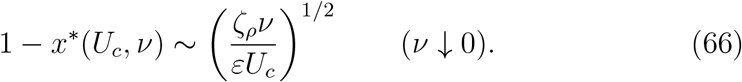

Positive repair therefore regularizes the no-repair boundary collision through a square-root displacement of the equilibrium from the fully damaged state.

Part (ii) of Theorem 1 also locates exactly where the hypothesis 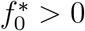 of Proposition 3 fails in the binary model: it holds for *U < U*_*c*_, while at *U* = *U*_*c*_ the interior equilibrium reaches the fully damaged boundary and for *U > U*_*c*_ no equilibrium with 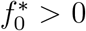 remains. The threshold *U*_*c*_ = 1 − *ζ*_*ρ*_*/ζ*_*γ*_ depends only on the fitness landscape, not on the demographic ratio *ε*.

The exact binary result also clarifies the relationship between the rare-mutation result and finite mutation pressure. The factor *λ/*(*λ* + *ν*) is not introduced by an independent approximation: it is the first-order limit of the exact nonlinear homeostatic equilibrium. Repair further changes the global equilibrium structure. Without repair, sufficiently strong mutational input drives the binary system to the fully damaged boundary, where the load saturates at *λ*. This is a transition in class composition, not extinction: total population size remains fixed by homeostatic regulation. Positive repair continually returns individuals to the pristine class and thereby maintains an interior balance for every 0 *< U* ≤ 1.

The exact binary solution also permits the equilibrium load to be followed across the full range of mutation pressure. Figure 1 shows this dependence for several repair rates.

**Figure 1:**
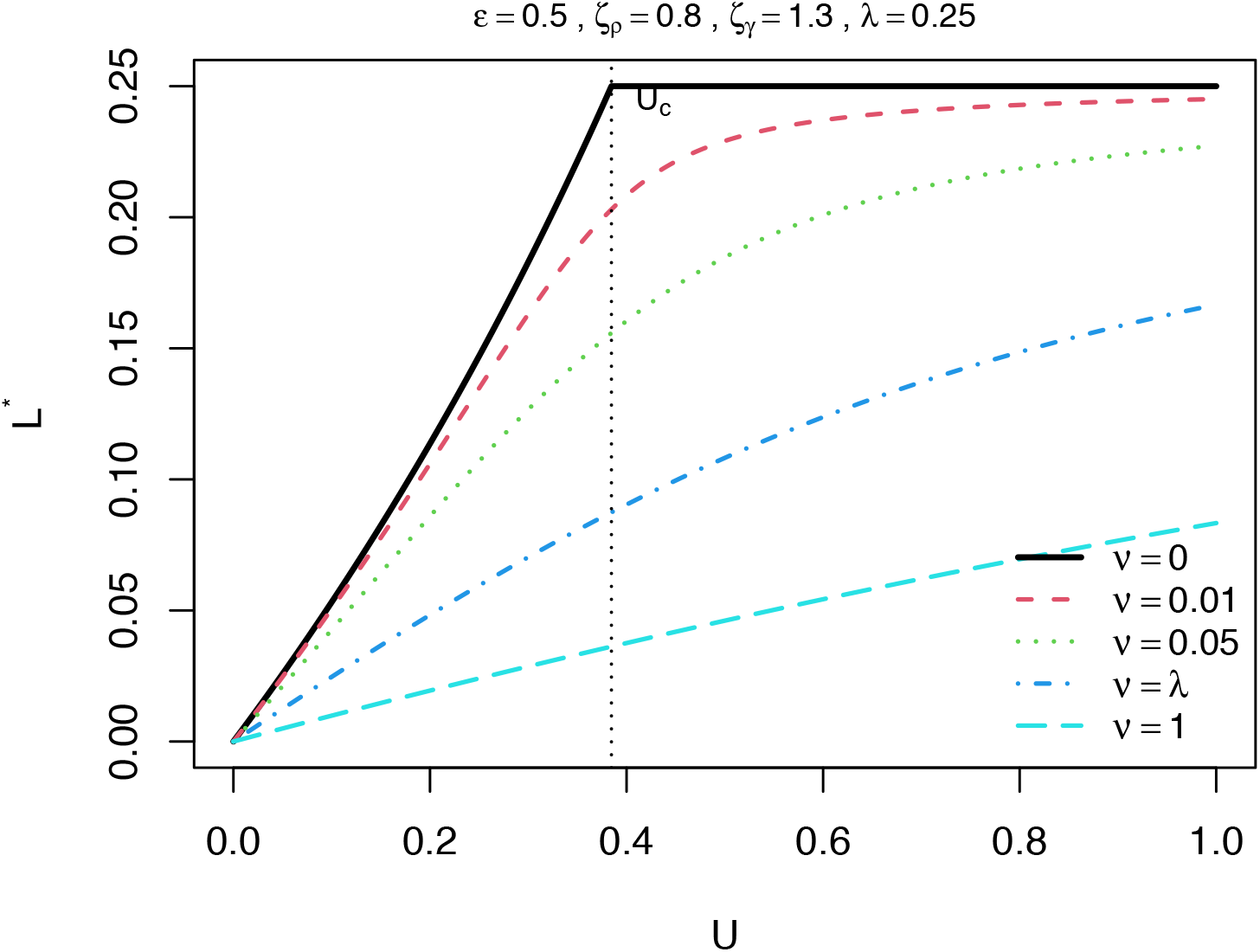
Exact binary equilibrium load *L*^∗^ = *λx*^∗^ as a function of mutation probability *U* for several repair rates *ν*. Parameters are *ε* = 0.5, *ζ*_*ρ*_ = 0.8, and *ζ*_*γ*_ = 1.3, giving *λ* = 0.25 and *U*_*c*_ = 1 − *ζ*_*ρ*_*/ζ*_*γ*_ ≈ 0.3846. In the absence of repair, the pristine-positive equilibrium branch reaches the fully damaged boundary at *U* = *U*_*c*_, where *L*^∗^ = *λ*. Positive repair removes this binary boundary equilibrium and yields an interior equilibrium across the full mutation range, with equilibrium load decreasing as the repair rate increases. The vertical dotted line marks the no-repair threshold *U*_*c*_.

### 3.6. Numerical validation of the asymptotic and exact results

Each asymptotic prediction was supported by the numerical calculations. For the full *n* = 10 multiclass system, with

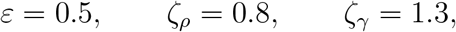

we have

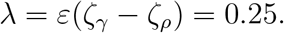

Repair was restricted to the first damaged class, with

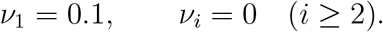

For panel A, multiclass equilibria were obtained by numerical integration of Eqs. (8)–(10) from the pristine state **e**_0_ using the LSODA solver implemented in R package deSolve, with relative and absolute tolerances 10^−11^ and 10^−13^, respectively. Integration proceeded in successive 100-unit intervals in *τ*, up to a maximum horizon *τ* = 5000, and was terminated once max_*i*_ |*df*_*i*_*/dτ* | *<* 10^−11^ and 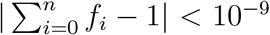. As an additional stability check, along the closed-form no-repair branch of Proposition 3 with *n* = 10, the Jacobian restricted to the simplex tangent space had strictly negative spectrum over the numerical sweep of 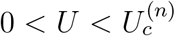. For example, the largest transverse eigenvalue was −0.2426 at *U* = 0.05 and −0.0997 at *U* = 0.9. Thus the branch is numerically linearly stable well beyond the small-*U* regime covered by Proposition 2, although no analytical stability result over the full branch is claimed here. For panel B, the binary equation (19) was integrated directly at *U* = *U*_*c*_ with *ν*= 0, initialized at *x*(0) = 0, and run to *τ* = 10^3^ with the same solver and tolerances; panel C evaluates the exact binary equilibrium root of Eq. (58) directly, without integration.

As shown in Fig. 2, the absolute error between the numerically computed equilibrium load and the leading-order repair-modified approximation,

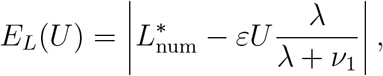

was approximately quadratic in *U*. A least-squares fit of log *E*_*L*_ against log *U* over the *U*-values shown in Fig. 2A gave a slope of 2.006, in close agreement with the *O*(*U* ^2^) remainder predicted by Corollary 1. Fitting log *E*_*L*_ against log *U* over 0.001 ≤ *U* ≤ 0.020, quadratic convergence was also obtained when repair was applied at every damaged class, with fitted slopes 2.0038 for *ν*_*i*_ = 0.1 and 2.0024 for *ν*_*i*_ = 0.5 for all *i* ≥ 1.

**Figure 2:**
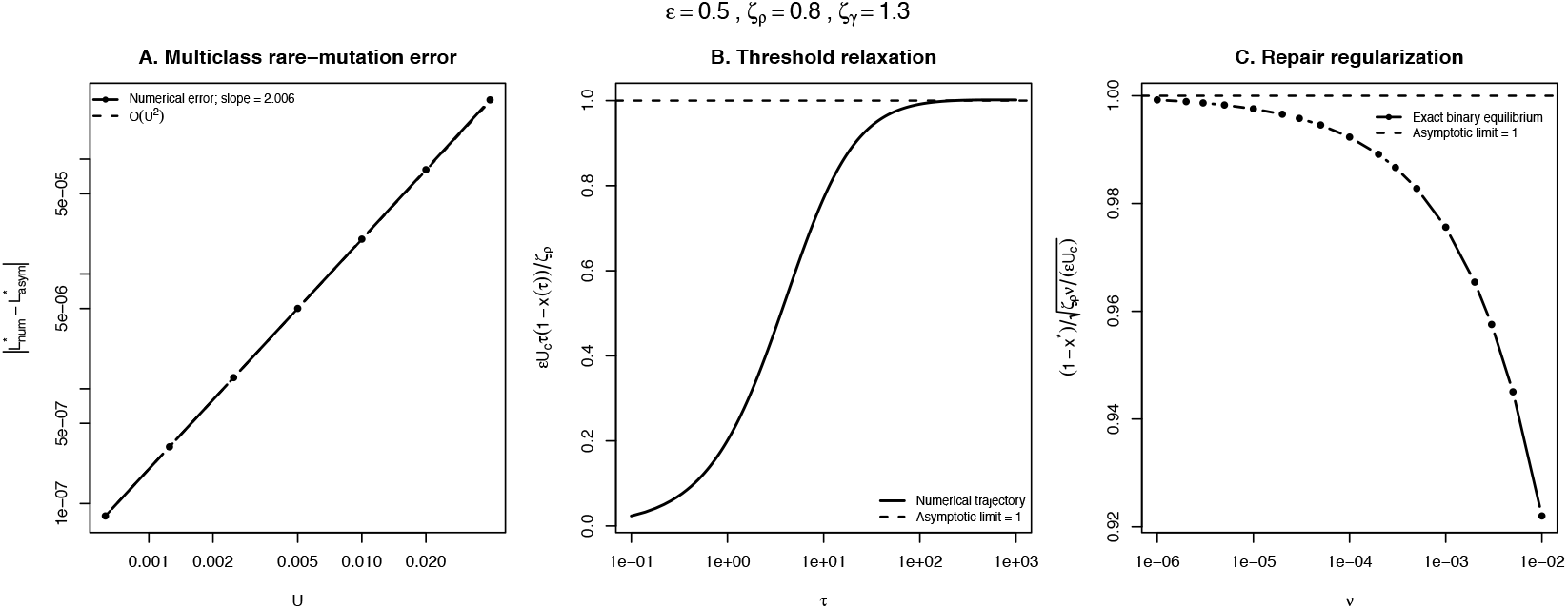
Numerical validation of the asymptotic and exact results. Common parameters are *ε* = 0.5, *ζ*_*ρ*_ = 0.8, and *ζ*_*γ*_ = 1.3, giving *λ* = *ε*(*ζ*_*γ*_ − *ζ*_*ρ*_) = 0.25 and, for the binary model, *U*_*c*_ = 1 − *ζ*_*ρ*_*/ζ*_*γ*_ = 0.384615 …. **(A)** Multiclass rare-mutation convergence for *n* = 10, with *ν*_1_ = 0.1 and *ν*_*i*_ = 0 for *i* ≥ 2. The absolute error 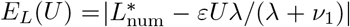 scales empirically as *U* ^2.006^ over the seven fitted values spanning 6.25*×*10^−4^ ≤ *U* ≤ 0.04, in agreement with the *O*(*U* ^2^) remainder in Corollary 1; the dashed line shows *U* ^2^ scaling. **(B)** Algebraic relaxation at the no-repair threshold *U* = *U*_*c*_. The normalized quantity *εU*_*c*_*τ* [1 − *x*(*τ*)]*/ζ*_*ρ*_ approaches 1, consistent with 1 − *x*(*τ*) ~ *ζ*_*ρ*_*/*(*εU*_*c*_*τ*); the dashed line denotes the asymptotic limit. **(C)** Repair regularization at *U* = *U*_*c*_. The exact binary equilibrium satisfies 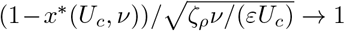 as *ν*↓ 0, consistent with the predicted square-root displacement from the fully damaged boundary. The dashed line denotes the asymptotic limit.

For the binary model, the same *ε, ζ*_*ρ*_, *ζ*_*γ*_ values give

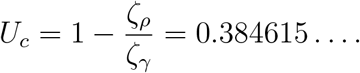

At the no-repair threshold *U* = *U*_*c*_, the normalized quantity

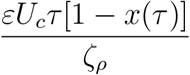

was 1.0015 at *τ* = 10^3^, consistent with the asymptotic prediction

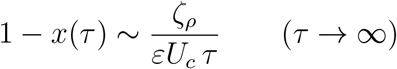

from Eq. (65).

Finally, at *U* = *U*_*c*_ with positive repair, the normalized equilibrium displacement

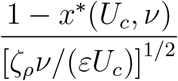

approached 0.9992 at *ν*= 10^−6^, the smallest repair rate considered. This agrees with the square-root regularization

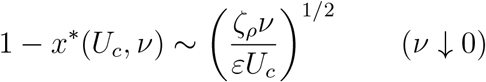

derived in Eq. (66). Together, these calculations provide numerical checks of the predicted *O*(*U* ^2^) multiclass remainder, algebraic threshold relaxation, and square-root regularization by repair.

#### Terminal repair and the multiclass boundary state

The binary regularization in Theorem 1 does not extend automatically to the multiclass model. In the binary system the damaged class is also the terminal class, so any *ν>* 0 provides an outward repair flux from the fully damaged state and removes *e*_1_ as an equilibrium. For *n >* 1, repair restricted to lower damage classes need not do so. In particular, if *ν*_*n*_ = 0, then *e*_*n*_ remains an exact equilibrium for every *U*, because no transition removes occupancy from the terminal class.

Thus any multiclass extension of the statement that positive repair removes the fully damaged boundary equilibrium must at minimum require *ν*_*n*_ *>* 0. More generally, when repair acts at every damaged class, the fully damaged boundary state is no longer an equilibrium. In the numerical checks reported here, trajectories initialized at both *e*_0_ and *e*_*n*_ converged to the same interior equilibrium when *ν*_*i*_ *>* 0 for all *i* ≥ 1. This behavior is consistent with multiclass regularization, but uniqueness and global attraction of an interior equilibrium for arbitrary *n* are not established here.

## 4. Discussion

Taken together, the results above have a common demographic interpretation, which we now develop and place against existing mutation–selection theory.

### Homeostatic mutation input and Haldane’s cancellation

The exact no-repair equilibrium identity, Eq. (53), shows that load equals *U* times realized reproductive turnover. In the uncapped hierarchy, this quantity is precisely the realized upward mutational flux, so equilibrium load equals realized deleterious input exactly for every 0 ≤ *U <* 1. Near the pristine state, realized turnover is *ε* + *O*(*U*), reducing this exact identity to

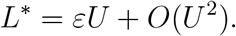

The factor *ε* in the unrepaired result has a direct demographic interpretation. The mutation probability *U* is specified per reproductive event, whereas time is measured in units of the pristine reproductive rate *ρ*_0_. At pristine homeostasis, births balance deaths and occur at dimensionless rate *ε* = *γ*_0_*/ρ*_0_.

In the no-repair rare-mutation limit, homeostatic regulation preserves the Haldane-type cancellation while converting mutation probability per reproductive event into the realized mutation input generated by demographic turnover. Along the pristine-positive no-repair branch, the exact relation

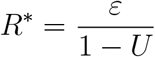

describes the reproductive adjustment required to sustain that balance.

This demographic interpretation echoes the longstanding observation that density regulation and excess reproductive production can buffer the effect of genetic deaths on population abundance [15]. The present model gives that qualitative idea a precise dynamical form: load is not inferred solely from the number of genetic deaths, but is represented exactly by the additional reproductive effort required to maintain homeostasis.

Proposition 1 strengthens this interpretation beyond the rare-mutation equilibrium. Under the definition of load used here,

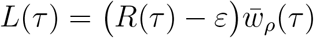

for every admissible state. Load is therefore exactly the excess reproductive effort above the pristine mortality-matched baseline, weighted by the population’s current mean fecundity. Repair does not enter this identity directly because its class-transition fluxes conserve population size. It changes load indirectly by redistributing individuals among classes and thereby changing both *R* and 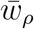.

### Repair partitions damaged-state removal

Repair modifies the Haldane-type cancellation not by changing the intrinsic fitness deficit of the damaged state, but by changing the selective share of first-order removal from that state. When *ν*≫ *λ*, restoration rather than selective depletion dominates that removal.

### Exact finite-U behavior

The binary model extends this interpretation beyond the rare-mutation limit. For *ν>* 0, the exact equilibrium is the unique biologically admissible root of the quadratic derived in Theorem 1, and its load is

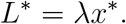

Its small-*U* expansion recovers *εUλ/*(*λ* + *ν*), establishing Corollary 1 as the local limit.

The binary model also reveals a qualitative effect that is invisible in the leading-order expansion. Without repair, the interior equilibrium reaches the fully damaged boundary at the no-repair collision *U* = *U*_*c*_ of Eq. (61). For *U > U*_*c*_, the attracting biological equilibrium is *x* = 1. The collision also has a distinctive dynamical signature. At *U* = *U*_*c*_ and *ν*= 0, the boundary equilibrium is nonhyperbolic and is approached algebraically rather than exponentially:

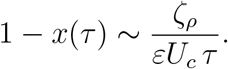

Thus the threshold is accompanied by slow algebraic relaxation in composition, even though total population size remains fixed.

Positive repair unfolds this degeneracy. At the collision value *U* = *U*_*c*_, the repaired equilibrium remains interior and satisfies

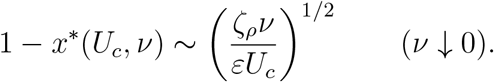

The square-root displacement shows that repair regularizes the no-repair boundary collision singularly rather than by a linear shift of the equilibrium. Thus the binary threshold is a transition in class composition, not population extinction. Any positive repair rate removes the boundary collision for *U >* 0 by continually returning individuals to the pristine class and maintaining an interior equilibrium.

The boundary transition in Theorem 1 is related to, but should not be identified uncritically with, the error thresholds of quasispecies and mutation– selection theory [7, 8, 9, 10]. In that literature, an error threshold generally denotes the loss or delocalization of a favored genotype under sufficiently strong mutation, and its existence and form can depend sensitively on the fitness landscape and mutation model. The transition derived here is instead the exact boundary collision of a finite two-class homeostatic system at the collision value *U*_*c*_ derived above. It is therefore not proposed as a universal error threshold. Its specific features are that population abundance remains fixed, the load saturates at the full damaged-class deficit, and every positive repair rate restores an interior equilibrium for *U >* 0.

The dependence of the multiclass boundary collision on terminal closure can be characterized exactly. In the uncapped no-repair hierarchy, the corresponding pristine-positive equilibrium remains normalizable for every 0 ≤ *U <* 1. Consistently, the finite-class endpoints of Proposition 3 satisfy

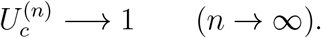

The finite-*n* collision is therefore a consequence of truncating the damage hierarchy rather than a universal mutation threshold of the uncapped model. The leading rare-mutation result *L*^∗^ = *εU* + *O*(*U* ^2^) is independent of this distinction.

### Relation to Muller’s ratchet

The loss of the least-damaged class is also the defining event in Muller’s ratchet, in which finite asexual populations can stochastically lose their least-loaded class through drift [22, 23]. The mechanism here is distinct. In the finite deterministic hierarchy, the pristine-positive branch reaches the boundary at 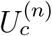, whereas in the uncapped deterministic hierarchy the corresponding equilibrium retains 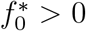 for every *U <* 1. In a finite-population extension, one would therefore expect stochastic loss of the pristine class to become important when its expected occupancy 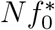 becomes small, even below any deterministic boundary collision. Repair provides a separate restoration flux toward lower-damage classes and would consequently oppose such irreversible loss, paralleling the role of reverse or compensatory transitions in stochastic ratchet models.

### Landscape dependence enters locally at first order

Without repair, the class-0 to class-1 fitness contrast cancels from the leading load. With repair, it remains in

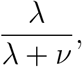

so the load increases with *λ* when *ν*is held fixed. Stronger selective differences reduce the frequency of the damaged class, but they also increase the fraction of that class removed through selection before repair can restore it.

This dependence should be interpreted narrowly. At order *U*, only the first damaged class contributes, so the fitness landscape enters through the single local coefficient

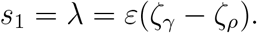

Higher-class coefficients, repair rates, and interactions among multiple damaged classes contribute only at order *U* ^2^ or beyond. Repair thus changes the leading load from independent of *s*_1_ to dependent on *s*_1_; genuinely multiclass landscape dependence is a higher-order effect.

### Demographic limits

The rare-mutation slope makes the interaction between turnover and repair explicit:

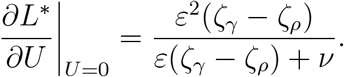

When *ν*= 0, this reduces to *ε*; equivalently, since 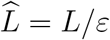,

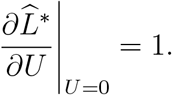

For fixed *ν* > 0,

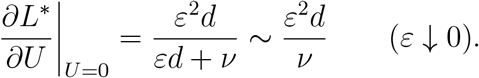

Thus repair changes the low-turnover scaling from linear in *ε* when *ν*= 0 to quadratic in *ε* when repair remains fixed. The crossover between selection-dominated and repair-dominated first-class removal occurs on the scale

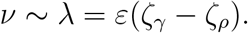

### Relation to previous mutation–selection theory

Hermisson et al. [7] analyze deterministic haploid mutation–selection systems generated by a fixed linear mutation–selection operator, with the nonlinear frequency dynamics arising through subtraction of scalar mean fitness. Their ancestral-distribution and maximum-principle results exploit the spectral structure of that fixed operator. The present system is not a special case of this formulation. Here the reproductive and mutational terms are multiplied by the state-dependent feedback

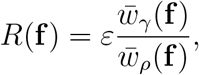

whose value depends nonlinearly on the full class distribution, while the mortality and repair operators are not multiplied by the same factor. Consequently, the dynamics cannot be represented by a fixed mutation–selection operator followed only by scalar mean-fitness normalization, and the maximum principle of Hermisson et al. does not directly apply. Likewise, neither the repair-induced regularization of the binary boundary collision nor the exact demographic identity of Proposition 1 follows as a direct specialization of their fixed-operator analysis.

Recent work by Pollett [17] provides a closer demographic comparison. In the deterministic large-population limit of that model, two-type mutation– selection dynamics are coupled to density-dependent population regulation, so total population density remains dynamic and mutation contributes an effective mortality that helps determine whether the deterministic system admits a positive population equilibrium. Here, by contrast, total abundance is constrained exactly by homeostatic feedback, and the demographic effect of mutation is expressed through compensatory reproductive effort rather than population decline. Accordingly, the threshold *U*_*c*_ derived here is not a deterministic persistence threshold but a boundary collision in class composition; in the no-repair binary system, the load instead saturates at the full damaged-state deficit at and above the collision. Thus the two models address complementary consequences of demographic regulation in mutation– selection systems.

A complementary infinite-dimensional perspective is provided by Pollett [21], who studies mutation–selection balance on a countably infinite trait space under density-dependent regulation. There, whether a localized positive equilibrium exists depends on the tail behaviour of selection: a confining mortality profile, increasing without bound in the tails, makes the associated mutation–mortality operator compact and yields a spectral threshold for equilibrium, whereas bounded mortality may allow mutation to transport mass indefinitely through trait space, so that no stationary trait distribution need exist. Proposition 4 is consistent with that picture: the geometric mortality 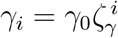 is confining along the relevant one-sided tail, and the un-capped hierarchy remains normalizable. It does not follow from that result, however: reproduction here is class dependent rather than trait independent, mutation is directional in the no-repair hierarchy rather than governed by an irreducible kernel on Z, and total abundance is held exactly constant by *R*(*f*) rather than regulated toward a population ceiling.

Backward transitions have also previously been included in general mutation– selection models [5, 7]. At leading order, the competing-exit structure found here is consistent with the familiar balance

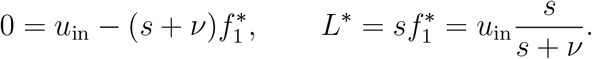

The present result identifies

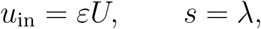

within an endogenously regulated demographic system. The factor *λ/*(*λ* + *ν*) should therefore not be understood as arising only under homeostasis. The specifically homeostatic contributions are that the realized mutation input is generated by demographic turnover, the feedback *R* depends nonlinearly on the full class distribution, and load has the exact demographic representation of Proposition 1. The exact binary solution further shows how these components interact at finite mutation pressure.

Wachter, Evans, and Steinsaltz [12] obtain an exact nonlinear generalization of Haldane’s principle in an age-structured demographic mutation– accumulation model. In their formulation, allelic outflow at mutation–selection balance is governed by marginal selective costs that depend on the genetic background, and their Haldane relation is an equilibrium balance between continuing mutational input and those marginal costs. The nonlinearity here has a different origin: it enters through the scalar feedback *R*(**f**), which rescales reproductive and mutational fluxes but not mortality or repair while holding total abundance exactly constant. Correspondingly, Proposition 1 identifies load with excess reproductive effort and, unlike the equilibrium Haldane relation of Wachter et al., is an algebraic identity valid at every admissible state.

More broadly, the present model differs from age-structured population-genetic and mutation-accumulation theories [11, 12, 24], in which mutations alter complete age-specific demographic schedules and selection is expressed through effects distributed across ages. Here the structure variable is accumulated damage, and population-level nonlinearity enters through exact regulation of total abundance.

### Scope and limitations

Repair is represented as a conservative downward transition between damage classes. Biologically, it may describe molecular repair, physiological restoration, epigenetic resetting, or another process that returns an individual toward a lower-damage state. Mathematically, it is analogous to backward mutation, although the modeled state is damage rather than DNA sequence. Repair is assumed to carry no direct reproductive, mortality, or energetic cost. Consequently, in the binary model the equilibrium load decreases monotonically with repair rate, as shown in Theorem 1. The binary model therefore does not address an optimal repair rate: within that model, without an opposing cost, increasing repair always reduces the load defined here. An interior optimum would require an explicit reproductive, mortality, energetic, or fidelity cost of repair, which would alter both the homeostatic feedback and the definition of total load.

Exact homeostasis also becomes increasingly demanding near the branch endpoint. Along the no-repair branch, *R*^∗^ = *ε/*(1 − *U*), so

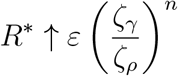

as 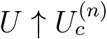 at the parameters of Fig. 2, with *n* = 10, this is approximately 128 times the pristine value. The feedback *R* is a reproductive multiplier rather than a hard bound, but biological systems have finite reproductive capacity, so an upper constraint on reproductive output could become important before this formal endpoint is reached. The strong-mutation behavior of the capped hierarchy should therefore be interpreted as a property of the exact-homeostasis idealization.

For the finite no-repair hierarchy, the equilibrium set can be classified explicitly. Above the pristine-positive branch endpoint, the terminal state **e**_*n*_ is the unique equilibrium and is locally asymptotically stable. A proof of global convergence to **e**_*n*_ for arbitrary *n*, however, is not pursued here. Proposition 4 concerns equilibrium existence and normalization only; well-posedness and dynamical stability of the corresponding infinite-dimensional system are not analyzed here.

The general multiclass result is a local result for *U* → 0 with fixed finite demographic and repair parameters. Joint limits in which repair rates vary with *U* require separate expansion. For example, if *ν*= *θU* with fixed *θ*, direct expansion of the first-class balance gives

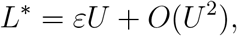

so repair first becomes distinguishable from the no-repair result at second order. The exact binary result removes the small-*U* restriction but does so only for a single terminal damaged class.

Further extensions include calculating the multiclass *O*(*U* ^2^) term, allowing mutation to produce multi-step damage increments, allowing repair to skip or restore multiple damage classes, incorporating costs and imperfect fidelity of repair, and treating finite populations in which drift interacts with selection and restoration.

### Conclusion

Homeostatic regulation links mutation load to demographic turnover, while repair changes the selective share of first-order damaged-state removal. Without repair, along the equilibrium branch with positive pristine-class frequency, load equals the mutation probability times realized per-capita reproductive turnover exactly, and in the uncapped hierarchy equals the realized deleterious input exactly. With repair, the leading-order load is reduced by the factor *λ/*(*λ* + *ν*). The exact demographic identity extends this interpretation beyond equilibrium, while the finite-*U* binary solution shows that the no-repair system undergoes a nonhyperbolic boundary collision with algebraic relaxation, that any positive repair rate removes this collision, and that, at the no-repair threshold, the repaired equilibrium is displaced from the boundary on a square-root scale as *ν*↓ 0. Together, these results distinguish the familiar competing-transition structure inherited from classical mutation–selection theory from the specifically homeostatic effects of turnover-dependent mutation input and nonlinear demographic feedback.

## Acknowledgments

The author thanks Fred Adler, Jon Seger, and Alan Rogers for many helpful discussions on population genetics and mathematical modeling. This research did not receive any specific grant from funding agencies in the public, commercial, or not-for-profit sectors.

## Code availability

The R script reproducing all numerical results and Figures 1 and 2 is available at GitHub: https://github.com/Priceeqn1/Rdiffeqnscripts/blob/main/Haldane1_and_2_combined_figureplotscript_v2.R.

## Declaration of Generative AI and AI-Assisted Technologies in the Manuscript Preparation Process

During the preparation of this work, the author used Claude (Anthropic) and ChatGPT (OpenAI) to assist with manuscript editing, language refinement, organizational suggestions, and discussion of mathematical exposition. The scientific ideas, mathematical derivations, model development, and conclusions are the author’s own. After using these tools, the author reviewed and edited all content and takes full responsibility for the content of the published article.

